# Dedifferentiation of caudate functional organization is linked to reduced D1 dopamine receptor availability and poorer memory function in aging

**DOI:** 10.1101/2024.03.18.585623

**Authors:** Saana M. Korkki, Jarkko Johansson, Kristin Nordin, Robin Pedersen, Lars Bäckman, Anna Rieckmann, Alireza Salami

**Affiliations:** Aging Research Center, Karolinska Institute and Stockholm University, Solna, Sweden; Department of Medical and Translational Biology, Umeå University, Umeå, Sweden; Department of Diagnostics and Intervention, Umeå University, Umeå, Sweden; Umeå Center for Functional Brain Imaging, Umeå University, Umeå, Sweden; Wallenberg Centre for Molecular Medicine, Umeå University, Umeå, Sweden; Department of Human Sciences, University of the Bundeswehr Munich, Germany

**Author notes:** Corresponding author: Saana M. Korkki.

**Keywords:** aging, dopamine, striatum, functional connectivity, neural dedifferentiation, memory

## Abstract

Age-related alterations in cortico-striatal function have been highlighted as an important determinant of declines in flexible, higher-order, cognition in older age. However, the mechanisms underlying such alterations remain poorly understood. Computational accounts propose age-related dopaminergic decreases to impoverish neural gain control, possibly contributing to reduced specificity of cortico-striatal circuits, that are modulated by dopamine, in older age. Using multi-modal neuroimaging data (fMRI, PET) from a large lifespan cohort (*n* = 180), we assessed the relationship between dopamine D1-like receptors (D1DRs) and cortico-striatal function during rest and an n-back working memory task. The results revealed gradual age-related decreases in the specificity of functional coupling between the centrolateral caudate and cortical association networks during both rest and working memory, which in turn was associated with poorer short and long-term memory performance with older age. Critically, reduced D1DR availability in the caudate and the prefrontal cortex predicted less differentiated caudate-cortical coupling across the lifespan, in part accounting for the age-related declines observed on this metric. These findings provide novel empirical evidence for a key role of dopamine in maintaining functional specialization of cortico-striatal circuits as individuals age, aligning with computational models that propose deficient catecholaminergic neuromodulation to underpin age-related dedifferentiation of brain function.

## 1. Introduction

Aging is associated with alterations in various functional, structural, and neurochemical properties of the brain that contribute to cognitive decline in older age (Grady, 2012; Jagust, 2013; Nyberg et al., 2012). On the functional level, a common finding in past neuroimaging studies has been decreased selectivity and specificity of brain function in older age, known as age-related *neural dedifferentiation* (reviewed in Koen et al., 2020; Koen & Rugg, 2019). During cognitive task performance, older adults display reduced stimulus- and process-specificity of regional brain activity in comparison to younger adults (Dennis & Cabeza, 2011; Park et al., 2004; Srokova et al., 2024; Trelle et al., 2019). Moreover, the organization of large-scale functional brain networks has been shown to become less segregated with advancing age (Chan et al., 2014; Geerligs et al., 2014, 2015; Pedersen et al., 2021), reflecting age-related decreases in functional connectivity between regions belonging to the same functional network, and increases in connectivity between regions belonging to different functional networks (Chan et al., 2014; Damoiseaux, 2017; Geerligs et al., 2015).

This reduced specificity of brain function has been shown to be associated with decreased cognitive performance in older age (Koen et al., 2020; Malagurski et al., 2020; Pedersen et al., 2021), however, the underlying neurobiological mechanisms remain unclear. Aging is associated with decreasing integrity of ascending neuromodulatory systems that act to tune the function of widespread cortical and subcortical circuits (Cools, 2019; Shine et al., 2021). In particular, the dopaminergic neurotransmitter system exhibits age-related deterioration (Johansson et al., 2023; Karalija et al., 2022; Karrer et al., 2017) that has been linked to decreases in higher-order cognition, including short- and long-term memory, in older age (Bäckman et al., 2000; Landau et al., 2009; Nyberg et al., 2016). Dopamine, and other neuromodulators, alter neuron’s responsivity to inputs, optimizing the signal-to-noise properties of neural networks (Servan-Schreiber et al., 1990; Shine et al., 2021). Long-standing computational accounts suggest age-related dopaminergic decreases to attenuate neural gain, resulting in less distinctive neural representations and networks in the aging brain (Li et al., 2000, 2001; Li & Rieckmann, 2014; Li & Sikström, 2002). However, while these accounts correspond to observations from functional neuroimaging studies indicating reduced specificity of brain function in older age, direct in vivo evidence linking markers of the dopaminergic system to measures of neural dedifferentiation in aging remains scarce.

In the human brain, dopaminergic modulation of striatal and prefrontal regions plays a crucial role in supporting flexible, goal-oriented behaviour (Cools, 2019; Cools & D’Esposito, 2011; Gerfen & Surmeier, 2011; Ranganath & Jacob, 2016; Westbrook et al., 2021). The striatum communicates with cortical regions through multiple topographically-organized cortico-striato-thalamo-cortical loops. Specifically, the caudate and anterior parts of the putamen couple with heteromodal cortical regions implicated in higher-order cognitive function, whereas the posterior putamen and ventral striatum connect with cortical regions implicated in motor and reward-related processes, respectively (Alexander et al., 1991; Haber & Knutson, 2010; Parent & Hazrati, 1995). Early animal research employing tract tracing techniques demonstrated projections to vary along the rostral-caudal axis (Kemp & Powell, 1970), which was subsequently linked to anterior-posterior frontal cortical regions (Haber, 2003). Human imaging studies have further provided evidence for more intricate organization of cortical connectivity within the caudate, with converging zones receiving projections from distinct heteromodal cortical regions (Choi et al., 2017; Jarbo & Verstynen, 2015). Notably, a medial-lateral gradient of caudate functional connectivity with cortical association networks has been reported (Choi et al., 2012; Kosakowski et al., 2024; O’Rawe & Leung, 2022; Rieckmann et al., 2018), with more centrolateral parts of the caudate preferentially coupling with the cortical fronto-parietal network (FPN) that supports task-general cognitive control, while a more medial wall zone integrates with the cortical default-mode network (DMN) implicated in internally-oriented cognition (Choi et al., 2012; Rieckmann et al., 2018)

Interestingly, previous work examining age-related differences in caudate functional organization has found aging to be associated with dedifferentiation of centrolateral caudate connectivity with these two cortical networks, driven by decreases in caudate-FPN connectivity and increases in caudate-DMN connectivity in older age (Rieckmann et al., 2018). However, it remains unclear whether such dedifferentiation may in part be related to age-related alterations in dopaminergic modulation of cortico-striatal circuits. No evidence for an association between striatal dopamine transporter (DAT) availability, which mediates the reuptake of dopamine from the synaptic cleft (Elsworth & Roth, 1997), and caudate-cortical connectivity was observed in this previous study (Rieckmann et al., 2018). Functional coupling within the cortico-striatal circuits may be more closely related to post-synaptic markers of the dopaminergic system, either in the striatum itself, or in the prefrontal cortex, which provides top-down control over striatal function (Ott & Nieder, 2019; Van Schouwenburg et al., 2012). Specifically, the activation of post-synaptic dopamine D1-like receptors (D1DRs) is proposed to enhance the stability of network dynamics (Durstewitz et al., 2000), and has previously been linked to functional connectivity within both cortico-cortical (Pedersen et al., 2024; Rieckmann et al., 2011; Roffman et al., 2016) and cortico-striatal (Johansson et al., 2023) networks.

Additionally, it remains unclear whether the pattern of age-related dedifferentiation of caudate-cortical connectivity previously observed during rest (Rieckmann et al., 2018) is similarly expressed during cognitive tasks taxing striatal and fronto-parietal circuits. Prior work has reported age-related decreases in the segregation of functional networks during both rest and task engagement (Chan et al., 2014; Geerligs et al., 2014, 2015; Pedersen et al., 2021; Raykov et al., 2024; Zhang et al., 2021). Moreover, older age has been shown to be associated with reduced ability to flexibly modulate functional networks to support cognitive task performance (Avelar-Pereira et al., 2017; Heinzel et al., 2017; Lugtmeijer et al., 2023). Integration of striatal and fronto-parietal regions is particularly critical for working memory, during which the striatum gates updating of memory representations maintained in cortical regions (D’Esposito & Postle, 2015; Frank et al., 2001; Murty et al., 2011). Indeed, prior work indicates functional coupling between the striatum and fronto-parietal regions to be enhanced with increasing working memory load (Salami et al., 2018). In contrast, decoupling of the cortical FPN and DMN is often seen during externally-oriented cognitive tasks, including working memory (e.g., Roffman et al., 2016; Sambataro et al., 2010, but see Vatansever et al., 2015). Additionally, task demands may influence the involvement of dopaminergic systems (Salami et al., 2019), with prior evidence suggesting differential involvement of striatal and extrastriatal D1DRs in modulation of cortical networks during rest and working memory (Roffman et al., 2016).

Here, we leveraged data from the largest human positron emission tomography (PET) study (*n* = 180) on D1DRs to date to investigate the role of dopaminergic decreases in age-related dedifferentiation of cortico-striatal function. Healthy adult volunteers (20-79 years old) underwent fMRI scans during rest and while performing an n-back working memory task with three load conditions (1-back, 2-back, 3-back). PET assessment of D1DR availability was conducted with the radioligand [^11^C]SCH2339. Specifically, we focused on D1DR availability in the caudate, the most age-sensitive dopamine-rich region in the current sample (Johansson et al., 2023), to assess the influence of local D1DR differences on caudate’s functional organization in terms of connectivity with the associative cortex (Choi et al., 2012; O’Rawe & Leung, 2022; Rieckmann et al., 2018). Moreover, we examined D1DRs in the prefrontal cortex, a region with high densities of D1DRs (Froudist-Walsh et al., 2023; Hall et al., 1994), to evaluate their potential involvement in top-down regulation of caudate-cortical connectivity during task execution. Based on prior work (Rieckmann et al., 2018), we expected older age to be associated with reduced specificity of centrolateral caudate connectivity with cortical association networks, as well as a decreased ability to modulate caudate-cortical connections in a task-dependent manner. Aligning with dopaminergic accounts of age-related neural dedifferentiation (Li et al., 2001; Li & Rieckmann, 2014; Li & Sikström, 2002), we further predicted reduced D1DR availability in the caudate and the prefrontal cortex to be associated with less differentiated caudate-cortical connectivity across the lifespan. Lastly, we expected less differentiated caudate-cortical connectivity during both rest and task to be associated with poorer performance on memory measures reliant on executive control operations supported by these networks.

## 2. Methods

The current study used baseline data from the DopamiNe, Age, connectoMe, and Cognition (DyNAMiC) cohort, described in detail in Nordin et al. (2022). Here, we report only methodological details relevant for the current work. Data collection for the DyNAMiC study was approved by the Regional Ethical board and the local Radiation Safety Committee of Umeå, Sweden.

### 2.1 Participants

The DyNAMiC sample consisted of 180 healthy adult volunteers, evenly distributed across the age range of 20 – 79 years old (mean age = 49.81, *SD*: 17.43; 50% female). Individuals were invited to participate via random selection from the population registry at Umeå, Sweden. Exclusion criteria included impaired cognitive function (Mini Mental State Examination score < 26), medical conditions or treatment that could affect brain or cognitive function (e.g., neurological, psychiatric, or developmental disorder, use of psychoactive medications, substance abuse, brain injury) or preclude participation in the neuroimaging assessments (e.g., metal implants). Individuals with other chronic or serious medical conditions (e.g., cancer, diabetes) were also excluded. All participants were right-handed native Swedish speakers and provided informed written consent prior to participation.

PET data on D1DR availability were missing from four participants due to drop out, technical issues, or indications of subcutaneous tracer injection, and caudate D1DR data were excluded for an additional three individuals due to unreliable D1DR estimates. Behavioural data for one or more of the memory tasks performed outside of the scanner were missing for 17 individuals due to technical issues or misunderstanding of task instructions. Outliers > 3.29 *SD*s from the mean were excluded from analyses of neuroimaging measures and cognitive tasks performed outside of the scanner. For the in-scanner n-back task, we restricted analyses to individuals who performed the task above chance-level accuracy (> 50%) for all load conditions (*n* = 165). Individuals with excessive movement in the scanner (mean framewise displacement; FD, > 0.30 mm, Jenkinson et al., 2002) were further excluded from analyses of the fMRI data (*n* = 1 for rest, *n* = 3 for n-back).

### 2.2 Image acquisition

MRI and PET scanning were performed at the Umeå Center for Functional Brain Imaging (UFBI) and the Umeå University Hospital in Umeå, Sweden.

#### 2.2.1 MRI

MRI scanning was performed with a 3T Discovery MR 750 scanner (General Electric) using a 32-channel phased-array head coil. High-resolution anatomical T1-weighted images were acquired with a 3D fast spoiled gradient-echo sequence (176 sagittal slices, slice thickness = 1mm, repetition time (TR) = 8.2 ms, echo-time (TE) = 3.2 ms, flip angle = 12°, field of view (FOV) = 250 × 250 mm, voxel size = 0.49 x 0.49 x 1mm). Functional MRI data were acquired during rest and an n-back working memory task using a T2*-weighted single-shot echo-planar imaging (EPI) sequence. The resting-state scan consisted of 350 volumes and the n-back scan of 330 volumes, acquired as 37 transaxial slices (slice thickness = 3.4 mm, interslice gap = 0.5 mm, TR = 2 000 ms, TE = 30 ms, flip angle = 80°, FOV = 250 × 250 mm, voxel size = 1.95 x 1.95 x 3.9 mm). During the resting-state scan, participants were instructed to stay awake and focus on a white fixation cross presented on a black background in the centre of the screen. The n-back task consisted of three load conditions (1-back, 2-back, 3-back) that were performed in a blocked fashion, described in detail below.

#### 2.2.2 PET

PET scanning was performed during rest with a Discovery PET/CT 690 scanner (General Electric) using the radioligand [^11^C]SCH23390. Head movements were minimized with individually fitted thermoplastic masks attached to the bed surface. Prior to tracer injection, a low-dose CT scan (10 mA, 120 kV, 0.8s rotation time) was acquired for PET attenuation correction. An intravenous bolus injection of [^11^C]SCH23390 with target radioactivity of 350 MBq was administered at the start of a 60min dynamic PET scan (6 x 10 s, 6 x 20 s, 6 x 40 s, 9 x 60 s, 22 x 120 s frames). The average radioactivity dose administered to participants was 337 ± 27 MBq (range 205–391 MBq). Time-framed, attenuation-, scatter-, and decay-corrected PET images (47 slices, 25 cm field of view, 256 × 256-pixel transaxial images, voxel size = 0.977 × 0.977 × 3.27mm) were reconstructed using the manufacturer-supplied iterative VUE Point HD-SharpIR algorithm (6 iterations, 24 subsets, resolution-recovery).

### 2.3 Image preprocessing and analyses

#### 2.3.1 FMRI

Functional data were preprocessed using Statistical Parametric Mapping 12 (SPM12, www.fil.ion.ucl.ac.uk/spm) and the Data Processing & Analysis of Brain Imaging toolbox (DPABI, version 6.1; Yan et al., 2016, http://rfmri.org/DPABI). Functional images were corrected for differences in slice acquisition time, movement, and distortion using subject-specific field maps. Distortion correction was not applied to data from three participants due to technical issues with field map acquisition. The functional data were coregistered with corresponding anatomical images and underwent nuisance regression to attenuate the influence of non-neural sources of noise. Nuisance regressors included mean cerebrospinal fluid (CSF), white matter (WM), and global signals, Friston’s 24-parameter motion model (Friston et al., 1996), and a binary scrubbing regressor indicating volumes contaminated by movement (i.e., FD > 0.2mm, Jenkinson et al., 2002). Data were bandpass filtered (0.009 – 0.09 Hz), normalized into MNI space using a sample-specific structural template created with Diffeomorphic Anatomical Registration Through Exponentiated Lie Algebra (DARTEL) toolbox (Ashburner, 2007), and spatially smoothed using an isotropic 6 mm full width at half maximum (FWHM) Gaussian kernel.

To characterize the topography of intrinsic caudate-cortical connectivity in the current dataset, we first conducted voxel-wise parcellations of resting-state connectivity between the caudate and the cortex in each age group (young, 20-39 years, *n* = 59; middle-aged, 40-59 years, *n* = 58; older, 60-79 years, *n* = 62), following the approach implemented in Choi et al. (2012). Specifically, for each individual, the correlation between each caudate voxel and each cortical voxel was computed. The correlation maps were averaged across participants within each age group, and each caudate voxel was then assigned to its most correlated cortical network, based on the network that was most frequently represented in the top 25 correlated cortical voxels. Cortical networks were defined based on the Yeo 7-network parcellation (Yeo et al., 2011), and the caudate based on the Harvard-Oxford subcortical atlas.

The voxel-wise parcellations unveiled a distinctive functional organization of the caudate along the medio-lateral axis in younger adults, similar to previous reports (Choi et al., 2012; Rieckmann et al., 2018), where more lateral regions of the caudate preferentially coupled with the FPN and more medial regions with the DMN. Further analyses of the group-wise parcellations revealed age-related alterations in this functional specialization. Specifically, the caudate subregion preferentially coupling with the FPN exhibited diminished connectivity with the FPN and increased connectivity with the DMN with advancing age (see Figure 1).

**Figure 1.**
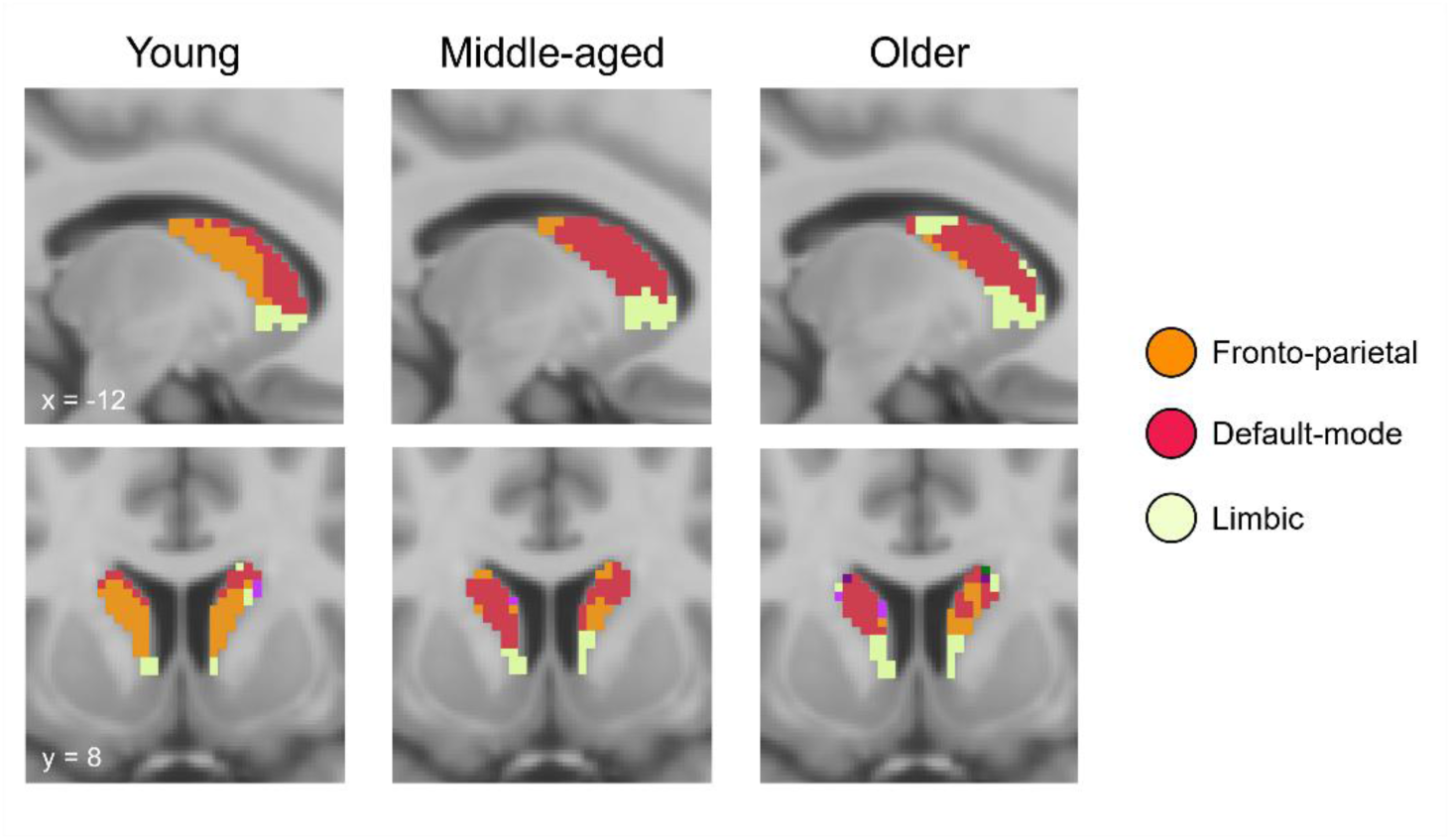
Voxel-wise parcellations of resting-state functional connectivity between the caudate and cortex in each age group. Caudate voxels are coloured based on the cortical network they were most strongly correlated with, based on a 7-network cortical parcellation (Yeo et al., 2011).

To facilitate quantification of individual differences in the strength of caudate functional connectivity with the FPN and the DMN, we followed the voxel-wise parcellations with region-of-interest (ROI) based analyses of functional connectivity. Specifically, we extracted time series from *a priori* ROIs, including a bilateral seed representative of the caudate FPN subregion (x, y, z = ± 12, 10, 8), and six bilateral cortical seeds representing three key nodes of the FPN (lateral prefrontal cortex, x, y, z = ± 41, 55, 4; medial prefrontal cortex, x, y, z = ± 5, 22, 47; anterior parietal cortex, x, y, z = ± 52, -50, 49) and three key nodes of the DMN (medial prefrontal cortex, x, y, z = ± 7, 46, -2; posterior parietal cortex, x, y, z = ± 42, -61, 31; precuneus, x, y, z = ± 3, -49, 25), consistent with the ROI definitions reported in Choi et al. (2012) and Rieckmann et al. (2018). Each ROI was created as a bilateral 6 mm radius sphere centred on the peak coordinates reported above.

Functional connectivity of the caudate seed to each of the two cortical networks (FPN, DMN) was computed as the average pairwise correlation (transformed to Fisher’s z) between the caudate seed and the cortical targets. For the n-back scan, connectivity was calculated separately for timepoints corresponding to each load condition (1-back, 2-back, 3-back), accounting for the hemodynamic lag (i.e., shifting timepoints by 4s, cf., Salami et al., 2018).

#### 2.3.2 PET

PET data were corrected for head movement using frame-to-frame image coregistration, coregistered with T1-weighted structural images, and resliced to dimensions of the structural images (1 mm^3^ isotropic, 256 x 256 x 256) using SPM 12. The T1-weighted structural images were segmented using FreeSurfer 6.0 (Fischl et al., 2002, https://surfer.nmr.mgh.harvard.edu) to obtain ROIs for the PET analyses. D1DR availability in target regions was estimated as the binding potential relative to non-displaceable binding in a reference region (BP_ND_, Innis et al., 2007), using the cerebellum as the reference. The simplified reference-tissue model (SRTM) was used to model regional time-activity course (TAC) data (Lammertsma & Hume, 1996). Regional TAC data were adjusted for partial volume effects (PVE) using the symmetric geometric transfer matrix (SGTM) method implemented in FreeSurfer (Greve et al., 2016), and an estimated point-spread-function of 2.5 mm FWHM. ROIs for the PET analyses included the bilateral caudate and the bilateral prefrontal cortex (comprising the caudal and rostral middle frontal gyrus, inferior frontal gyrus pars orbitalis and pars triangularis, and the frontal pole, which exhibited similar age-related D1DR trajectories in the current dataset, see Johansson et al., 2023).

### 2.4 Cognitive tasks

#### 2.4.1 In-scanner n-back task

The working memory task performed in the MRI scanner was a numerical n-back task consisting of three load conditions (1-back, 2-back, 3-back) completed in a blocked manner (c.f., Nevalainen et al., 2015; Nordin et al., 2022). In each task block, a sequence of 10 single numbers was presented on the screen (stimulus duration: 1.5s, inter-stimulus interval, ISI: 0.5s) and participants were instructed to indicate whether each item presented matched the one *n* items back (i.e., 1-, 2-, or 3-back) in the sequence by pressing one of two adjacent buttons on a scanner-compatible button box using their index and middle finger. Load condition for each task block was indicated by a cue presented on the screen before the start of the block. Nine task blocks were completed for each task load, with the order of blocks randomized but kept constant across participants. Performance in the n-back task was measured as the percentage of correct responses for each load condition.

#### 2.4.2 Memory tasks outside of the scanner

In addition to the in-scanner working memory task, participants completed a battery of episodic and working memory tasks outside of the scanner (Nordin et al., 2022). Episodic memory was assessed with a word recall, a number-word recall, and an object-location recall task. In the word recall task, participants were presented with 16 concrete nouns that appeared one by one on the computer screen (stimulus duration: 1s, ISI: 1s). After presentation of the entire list, participants used the keyboard to type in as many words as they could remember from the preceding list in any order. Two blocks of the word recall task were completed (maximum score = 32). In the number-word recall task, participants were presented with pairs of two-digit numbers and concrete plural nouns (e.g., 46 dogs). Ten number-digit pairs were first sequentially presented (stimulus duration: 6s, ISI: 1s), after which each word reappeared on the screen in a randomized order and participants were instructed to recall the associated number by typing their responses with the keyboard. Two blocks of the number-word recall task were completed (maximum score = 20). In the object-location recall task, 12 objects were sequentially presented in different locations on a 6 x 6 square grid on the computer screen (stimulus duration: 8s, ISI: 1s). Following the encoding phase, all objects appeared next to the grid and participants’ task was to place them in their correct location in the grid using the computer mouse. Participants could place the objects in any order. Two blocks of the object location recall task were completed (maximum score = 24).

Working memory was assessed with a letter updating, a number updating, and a spatial updating task. In the letter updating task, a sequence of capital letters (A-D, stimulus duration: 1s, ISI: 0.5s) appeared on the screen and participants were instructed to try and keep in mind the three most recently presented letters. When prompted, participants were asked to type the last three letters using the computer keyboard. The task consisted of 16 trials of 7, 9, 11, or 13 letter sequences (4 trials per sequence length), presented in a random order (maximum score = 48). The number updating task was a columnized numerical 3-back task, where a single digit number (stimulus duration: 1s, ISI: 0.5s) appeared in one of three boxes present on the screen, in a sequence from left to right. Participants’ task was to judge whether the current number matched the one previously presented in the same box (i.e., three numbers before) by pressing one of two assigned keys on the keyboard. Four task blocks each consisting of presentation of 30 numbers were completed (maximum score = 108). In the spatial updating task, three 3 x 3 square grids were presented next to each other on the computer screen. At the beginning of each trial, a blue dot appeared in a random location in each grid for 4s. After this, an arrow appeared below each grid to indicate the direction to which participants should mentally move the object to by one step. The arrows appeared sequentially from left to right (stimulus duration: 2.5s, ISI: 0.5s), and twice below each grid (i.e., each object should be moved by two steps). Participants were then asked to indicate where in each grid the object had moved to using the computer mouse. Ten blocks of the spatial updating task were completed (maximum score = 30).

Performance in all tasks was measured as the number of correct answers. To generate a composite score of memory performance, we performed a principal component analysis across all three episodic and all three working memory measures (z-scored). The first principal component from this analysis accounted for 54.96% variance in the data with high loadings across all tasks (*r*s .64 - .80, see Supplementary material), and was used as a measure of general memory function.

### 2.5 Statistical analyses

Statistical analyses were conducted with R (version 4.3.2, R Core Team 2023) and JASP (version 0.18.1, JASP Team 2023). Age group (i.e., young, 20-39 years; middle-aged, 40-59 years; older, 60-79 years) differences in caudate functional connectivity during rest and working memory were analysed with mixed ANOVAs. Linear regression and linear mixed effects models were used to assess the relationship between functional connectivity and D1DR availability, and functional connectivity and memory, during rest and working memory, respectively. Linear mixed effects models were implemented with the R package lme4 (version 1.1-35), including a random effect of participant and the fixed effects of interest. *P*-values were estimated via the Satterthwaite’s degrees of freedom method implemented in the R package lmerTest (version 3.1-3). Age, sex, and in-scanner movement (mean FD, Jenkinson et al., 2002) were included as covariates in all connectivity-D1DR and connectivity-memory analyses. Analyses of connectivity-memory relationships additionally controlling for educational level are reported in the Supplementary material. We further investigated whether D1DR availability mediated the effects of age on functional connectivity using the R package mediation (version 4.5.0). Mediation analyses were controlled for age, sex, and mean FD, and bootstrapped, bias-corrected, and accelerated confidence intervals were estimated with 5000 samples.

## 3. Results

### 3.1 Voxel-wise parcellations of caudate-cortical connectivity

To assess the overall functional organization of the caudate, we first performed voxel-wise mapping of resting-state functional connectivity between the caudate and the cortex within each age group (young, 20-39 years; middle-aged, 40-59 years; older, 60-79 years, see Figure 1). Consistent with previous work using similar approaches (Choi et al., 2012; Rieckmann et al., 2018), we observed that, in younger adults, caudate voxels were predominantly allocated to the FPN (39.58%) and the DMN (23.85%). Moreover, in the current dataset, 31.76% of caudate voxels in younger adults were allocated to the limbic network. Other cortical networks were allocated to only a small percentage of caudate voxels (< 5 %) in any age group (see Supplementary material) and are therefore not discussed further.

Of the voxels allocated to the FPN in younger adults, only 29.11% were allocated to the FPN in the middle-aged, and 22.03% were allocated to the FPN in the older adults, with a large proportion of these voxels instead being allocated to the DMN in these two age groups (middle aged: 58.73%, older adults: 50.13%). In contrast, 64.29% and 61.76% of the caudate voxels allocated to the DMN in younger adults were still allocated to the DMN in the middle-aged and older adults, respectively. Similarly, the caudate subregion coupling with the limbic network was relatively well-preserved in the middle-aged and older adults, with 81.70% of limbic voxels in younger adults allocated to the limbic network in the middle-aged, and 62.15% in the older adults. Thus, consistent with previous work (Rieckmann et al., 2018), the voxel-wise parcellations suggest particular vulnerability of the caudate subregion preferentially coupling with the FPN to age-related alterations.

### 3.2 Region-of-interest analyses

#### 3.2.1 Age-related dedifferentiation of caudate-cortical connectivity during rest

To quantify age-related differences in the strength of centrolateral caudate connectivity with the cortical FPN and DMN, we further performed ROI-based analyses of functional connectivity. For functional connectivity during rest, a mixed ANOVA indicated a significant interaction between age group (young, middle-aged, old) and cortical network (FPN, DMN), *F*(2, 175) = 8.63, *p* < .001, *η_p_^2^* = .09 (Figure 2A). Consistent with the pattern expected based on prior work (Choi et al., 2012; Rieckmann et al., 2018), younger adults demonstrated significantly stronger functional connectivity between the centrolateral caudate and the cortical FPN than the cortical DMN, *t*(58) = 4.85, *p* < .001, *d* = .63. However, no such preferential connectivity was detected in the middle-aged (*p* = .131) or older adults (*p* = .254). As suggested by the voxel-wise parcellations, this age-related dedifferentiation of functional connectivity of the centrolateral caudate was driven both by age-related reductions in caudate-FPN connectivity (i.e., the preferred network in younger adults), *F*(2,175) = 6.00, *p* = .003, *η_p_^2^* = .06, and age-related increases in caudate-DMN connectivity (i.e., the non-preferred network in younger adults), *F*(2,175) = 5.61, *p* = .004, *η_p_^2^* = .06. Specifically, we observed caudate-FPN connectivity to be decreased in older in comparison to the young, *t*(119) = 2.68, *p* = .009, *d* = .49, and middle-aged adults, *t*(117) = 3.00, *p* = .003, *d* = .55, whereas elevated caudate-DMN connectivity was observed in both older, *t*(119) = 2.37, *p* = .020, *d* = .43, and middle-aged adults, *t*(114) = 3.39, *p* < .001, *d* = .63, when compared to the young adults.

**Figure 2.**
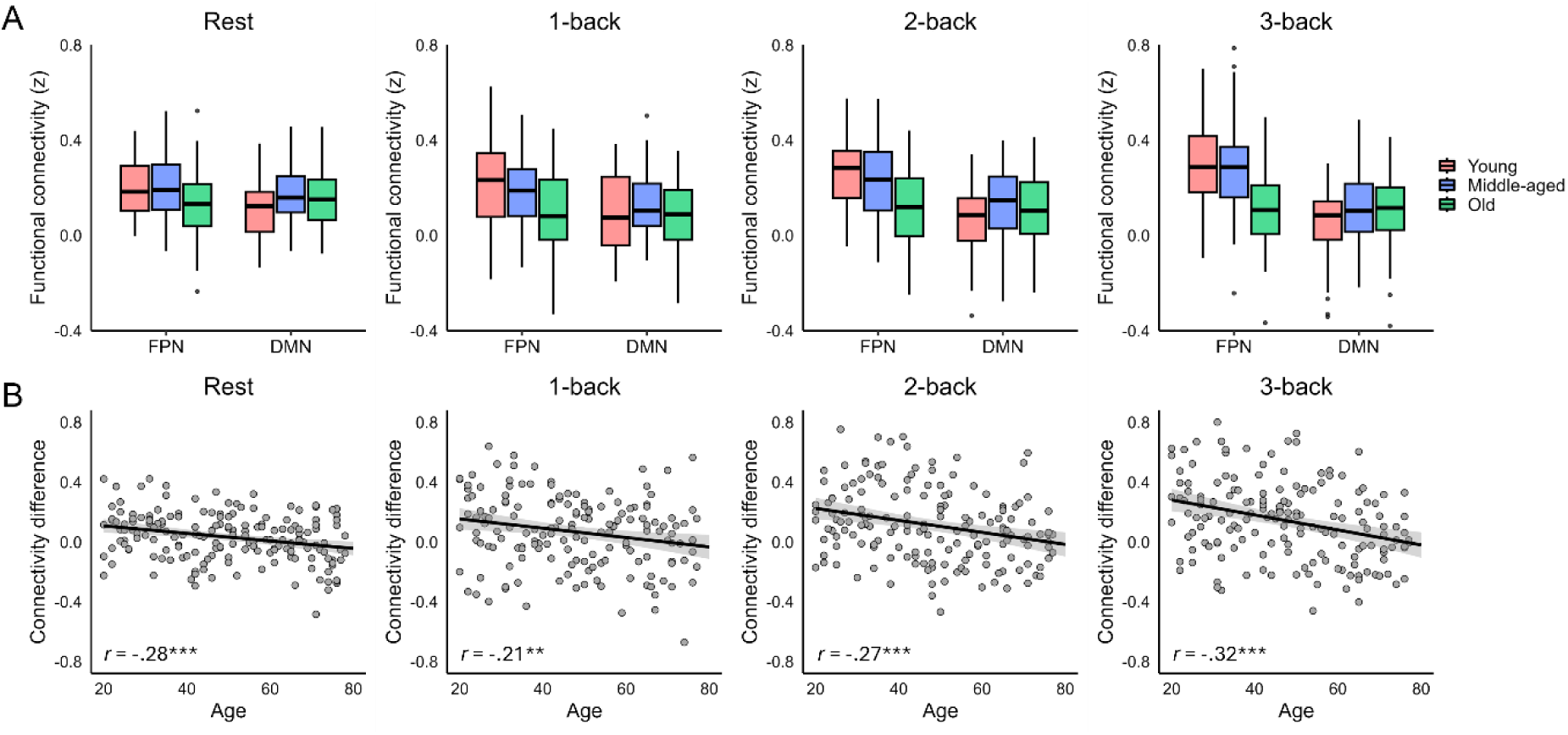
A) Functional connectivity (Fisher’s z) between the centrolateral caudate and the fronto-parietal (FPN) and default-mode (DMN) networks in each age group (young, 20-39 years, middle-aged, 40-59 years, old, 60-79 years old) and condition. B) Associations between age and the differentiation of caudate-cortical connectivity (i.e., the difference in caudate-FPN vs. caudate-DMN connectivity) in each condition.

#### 3.2.2 Age-related dedifferentiation of caudate-cortical connectivity during working memory

We next examined whether the age-related dedifferentiation of caudate-cortical connectivity observed during rest was similarly present during a working memory task that places greater demands on coordinated activity of striatal and frontoparietal regions (D’Esposito & Postle, 2015; Nyberg & Eriksson, 2016). During the n-back working memory task that participants performed in the scanner, we also observed a significant interaction between age group and cortical network, *F*(2, 160) = 10.13, *p* < .001, *η_p_^2^* = .11 (Figure 2A), but no further evidence for an interaction between age group and memory load (*p* = .810), or age group, memory load, and cortical network (*p* = .225). During n-back, both young, *F*(1, 58) = 48.49, *p* < .001, *η_p_^2^* = .46, and middle-aged adults, *F*(1, 56) = 21.42, *p* < .001, *η_p_^2^* = .28, exhibited stronger caudate-FPN over caudate-DMN connectivity across memory loads, whereas no significant differences between caudate-FPN and caudate-DMN connectivity were observed in the older adults (*p* = .511). Although the 3-way interaction between age group, memory load, and cortical network did not reach significance, we note that evidence for load-dependent upregulation of preferential caudate-FPN connectivity was observed in the young and middle-aged adults, as indicated by a significant interaction between load and cortical target in these age groups (young, *F*(2,116) = 3.60, *p* = .030, *η_p_^2^* = .06; middle-aged, *F*(2,112) = 4.66, *p* = .011, *η_p_^2^* = .08), but not in the older adults (*p* = .837). During working memory, age-related differences in functional connectivity were driven by reduced caudate-FPN connectivity in the older adults across memory loads (effect of age, *F*(2, 160) = 18.06, *p* < .001, *η_p_^2^ =* .18; old vs. young, *t*s > 3.35, *p*s < .002, old vs. middle-aged, *t*s > 2.78, *p*s < .007), whereas no significant age-related differences were detected for caudate-DMN connectivity during working memory (*p* = .127). Thus, older age was associated with dedifferentiation of caudate-cortical connectivity during both rest and working memory, with the age-related differences driven by both decreased caudate-FPN and increased caudate-DMN connectivity during rest and primarily by decreased caudate-FPN connectivity during working memory.

### 3.3 D1DR integrity contributes to maintaining caudate functional organization across the adult lifespan

Given the proposed role of dopamine in age-related dedifferentiation of brain function (Li et al., 2000, 2001; Li & Sikström, 2002) and in regulation of cortico-striatal connections (Cools, 2019; Gerfen & Surmeier, 2011), we next assessed whether integrity of the D1DR system contributes to maintaining the differentiation of caudate-cortical connectivity across the lifespan. For this purpose, we first quantified the degree of preferential caudate-FPN connectivity for each individual and task condition as the difference between centrolateral caudate connectivity with the FPN versus the DMN. The degree of preferential caudate-FPN connectivity was negatively associated with age in each task condition (*r*s -.21 to -.32, Figure 2B). Across the sample, caudate D1DR availability positively predicted the differentiation of caudate-cortical connectivity during rest, *β* = .22, *SE* = .10, *t* = 2.18, *p* = .031, controlling for age, sex, and mean FD (Figure 3A). This association did not significantly vary with age (*p* = .282). Prefrontal D1DR availability, on the other hand, was not significantly associated with the differentiation of caudate-cortical connectivity during rest (D1DR, *p* = .093, D1DR x age, *p* = .195), but predicted the differentiation of caudate-cortical connectivity during working memory, *β* = .14, *SE* = .07, *t* = 2.05, *p* = .042 (Figure 3B). The association between prefrontal D1DR availability and caudate-cortical connectivity did not significantly vary with memory load (*p* = .756), age (*p* = .493), or as an interaction between memory load and age (*p* = .722). Caudate D1DR was not significantly associated with the differentiation of caudate-cortical coupling during working memory (*p*s > .319). Given the different patterns of D1DR-connectivity relationships observed during rest and working memory, we further included both caudate and prefrontal D1DR in the same model to assess the specificity of these associations. The association between prefrontal D1DR and caudate-cortical connectivity during working memory persisted after inclusion of caudate D1DR availability as an additional covariate, *β* = .23, *SE* = .09, *t* = 2.46, *p* = .015, whereas the association between caudate D1DR and resting-state connectivity became marginally significant after inclusion of prefrontal D1DR availability, *β* = .23, *SE* = .13, *t* = 1.79, *p* = .075. This finding suggests that prefrontal D1DRs may exert a more pronounced influence on caudate-cortical coupling during a prefrontal-dependent task.

**Figure 3.**
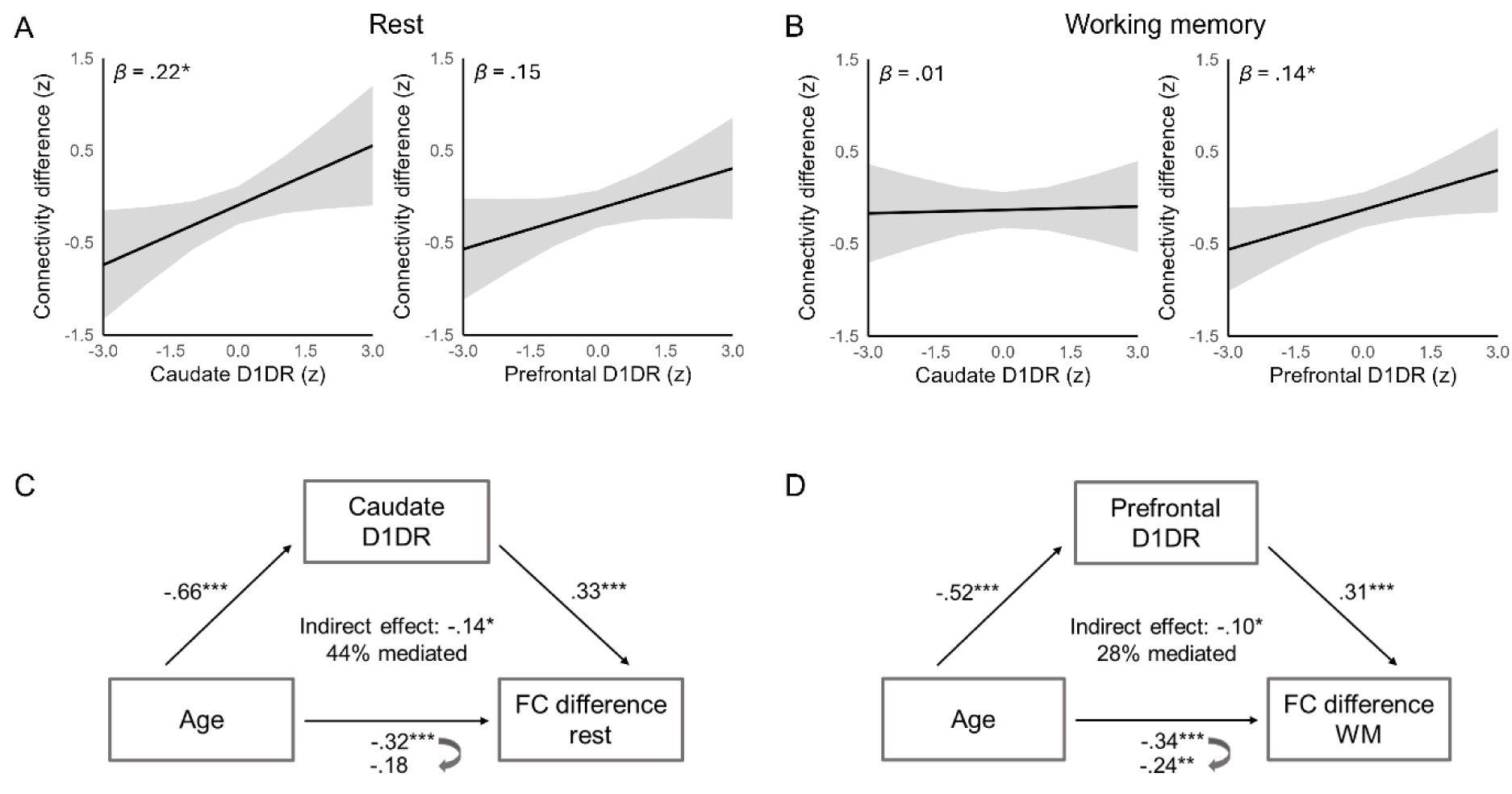
Associations between D1DR availability and the differentiation of caudate-cortical connectivity (difference in caudate-FPN and caudate-DMN connectivity) during A) rest and B) working memory. Plots illustrate predicted effects of D1DR availability on connectivity from A) linear regression and B) linear mixed effects analyses, controlling for A) age, sex, and mean framewise displacement (FD), and B) age, sex, mean FD, and working memory load. Mediation analyses indicated D1DR availability in the C) caudate and D) the prefrontal cortex to mediate the effect of age on the differentiation of caudate-cortical connectivity during C) rest and D) working memory, respectively.

To assess whether reduced D1DR availability accounted for age-related dedifferentiation of caudate-cortical connectivity, we further performed mediation analyses (Figure 3C and 3D). For functional connectivity during rest, we observed a significant indirect effect of age on the differentiation of caudate-cortical connectivity via caudate D1DR availability, *β* = -.14, 95% CI [-.24, -.04], *p* = .010, accounting for 44.14% of the age effect. The direct effect of age was not significant, *β* = -.18, 95% CI [-.38, .01], *p* = .072. Similarly, for the mean differentiation of caudate-cortical connectivity during working memory, we observed a significant indirect effect of age via prefrontal D1DR availability, *β* = -.10, 95% CI [-.21, -.01], *p* = .042, accounting for 28.35% of the age effect. For working memory, the direct effect of age was also significant, *β* = -.24, 95% CI [-.42, -.07], *p* = .008.

### 3.4 Dedifferentiation of caudate functional organization predicts poorer memory performance with older age

Lastly, we examined the behavioural relevance of differences in caudate-cortical coupling across the lifespan. We expected integrity of caudate-cortical circuits supporting flexible cognitive control to be associated with more efficient performance across a battery of short and long-term memory tasks. General memory function was indexed as the first principal component from an analysis involving all three episodic (i.e., word recall, number-word recall, object-location memory) and working memory (letter updating, number updating, spatial updating) tasks included in the cognitive task battery that participants performed outside the scanner (all task loadings *r*s > .64, see Supplementary material). Greater differentiation of caudate-cortical connectivity during rest was associated with better memory performance across the lifespan, *β* = .17, *SE* = .06, *t* = 2.80, *p* = .006, controlling for age, sex, and mean FD. We further observed a significant interaction between resting-state connectivity and age in predicting memory, *β* = .11, *SE* = .05, *t* = 2.06, *p* = .041, such that the association between connectivity and memory became stronger with advancing age (Figure 4A). Indeed, when examining associations between resting-state connectivity and memory within the three age groups separately, the differentiation of caudate-cortical coupling was positively associated with memory in the middle-aged, *β* = .37, SE = .13, *t* = 2.92, *p* = .005, and older individuals, *β* = .36, SE = .13, *t* = 2.73, *p* = .009, but not in the younger adults (*p* = .864). A similar interaction between age and resting-state connectivity was observed for the mean in-scanner n-back performance, *β* = .12, SE = .06, *t* = 2.02, *p* = .045.

**Figure 4.**
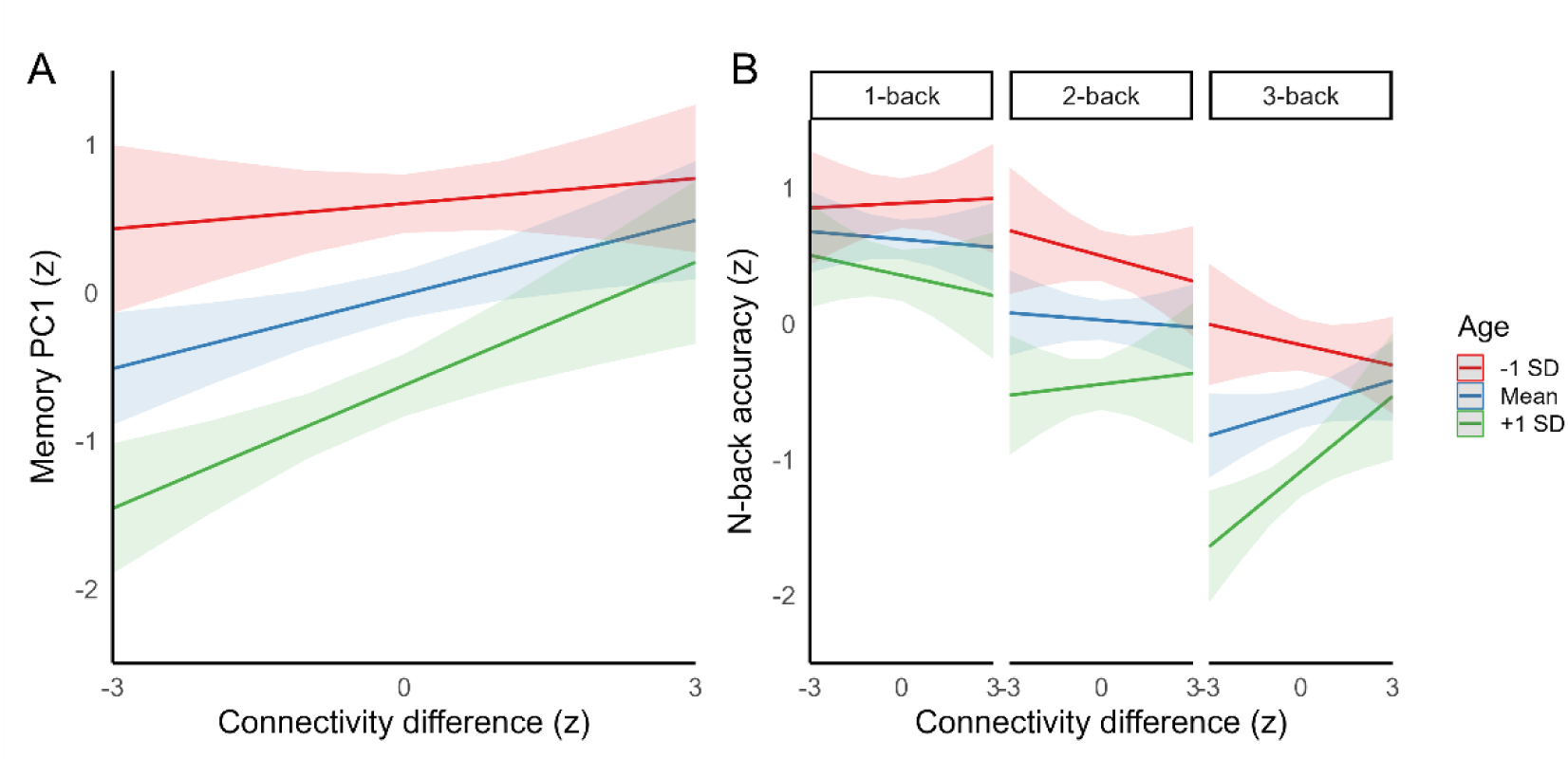
A) Association between the differentiation of caudate-cortical connectivity (difference between caudate-FPN and caudate-DMN connectivity) during rest and out-of-scanner memory performance (1^st^ principal component from an analysis including three episodic and three working memory tasks performed outside the scanner). B) Associations between the differentiation of caudate-cortical connectivity during working memory and in-scanner memory performance. Plots illustrate the predicted effects of connectivity on memory at mean ± 1 *SD* of age from A) linear regression and B) linear mixed effects analyses, controlling for age, sex, and mean framewise displacement (FD).

Assessing associations between functional connectivity during the n-back task and in-scanner working memory accuracy, we observed a significant 3-way interaction between age, memory load, and connectivity differentiation, *F*(2, 346.57) = 3.24, *p* = .040, where stronger preferential caudate-FPN connectivity was associated with better working memory performance with older age on the 3-back condition (see Figure 4B). Indeed, when examining associations between 3-back functional connectivity and 3-back accuracy within the age groups separately, we observed positive associations between the differentiation of caudate-cortical connectivity and working memory accuracy in the middle-aged, *β* = .27, SE = .13, *t* = 2.01, *p* = .049, and older adults, *β* = .29, SE = .14, *t* = 2.04, *p* = .048, but not in the younger group (*p* = .547). Differentiation of caudate-cortical connectivity during the in-scanner working memory task was not associated with performance on the memory tasks outside of the scanner (*p*s > .163). The interaction between age and resting-state connectivity in predicting memory, and the within-group associations between connectivity and memory for the in-scanner 3-back condition, did not survive inclusion of education as an additional covariate in the models (*p*s > .073). Otherwise, a similar pattern of results was observed when including educational level as an additional covariate (see Supplementary material).

## 4. Discussion

Using data from the largest human PET cohort on D1DRs to date, we examined the role of dopaminergic decreases in age-related alterations in cortico-striatal function. Consistent with previous findings (Rieckmann et al., 2018), advancing age was associated with less specific functional coupling between the centrolateral caudate and the cortical FPN. Such age-related differences were present during both rest and a working memory task that taxes coordinated activity of striatal and fronto-parietal regions (D’Esposito & Postle, 2015; Nyberg & Eriksson, 2016). Across the adult lifespan sample, lower D1DR availability in the caudate and the prefrontal cortex predicted less differentiated caudate-cortical coupling during rest and working memory, respectively, and partly accounted for age-related variation on this measure. Dedifferentiation of caudate-cortical connectivity was further associated with poorer memory function in midlife and older age. Together, these findings align with accounts proposing neural dedifferentiation as one mechanism underpinning cognitive decline in aging and highlight a critical contribution of the dopaminergic system to maintaining functional organization of cortico-striatal circuits across the lifespan.

Extending previous findings of particular vulnerability of the centrolateral caudate to age-related alterations in a sample of young and older individuals (Rieckmann et al., 2018), we here report gradual decline in the specificity of functional coupling between this caudate subregion and cortical association networks across the adult lifespan. Aligning with previous evidence indicating age-related decreases in functional network segregation during both rest and task states (Chan et al., 2014; Geerligs et al., 2014, 2015; Pedersen et al., 2021; Raykov et al., 2024; Zhang et al., 2021), this age-related dedifferentiation of caudate-cortical connectivity was consistently observed during both rest and while performing an n-back working memory task. During rest, age-related differences in functional specialization of the caudate were driven by both decreased caudate-FPN coupling (i.e., the preferred network in younger adults) and increased caudate-DMN coupling (i.e., the non-preferred network in younger adults) with older age. In contrast, during working memory, the age-related differences were primarily driven by decreases in caudate-FPN connectivity. Contrary to our expectations, age-related differences in caudate-cortical connectivity did not significantly vary with working memory load during the n-back task, although evidence for load-dependent modulation of caudate-cortical connectivity was limited to the young and middle-aged adults.

Importantly, the current study provides novel evidence for a role of decreased D1DR availability in age-related dedifferentiation of cortico-striatal function. This finding is consistent with prior studies linking PET markers of D1DR availability to functional coupling within cortico-cortical (Pedersen et al., 2024; Rieckmann et al., 2011; Roffman et al., 2016) and cortico-striatal (Johansson et al., 2023) circuits, and with pharmacological evidence showing dopamine depletion to reduce the stability of regional brain activity and the coherence of functional networks (Shafiei et al., 2019). At the single-neuron level, dopaminergic neuromodulation alters neuron’s sensitivity to inputs, enhancing the signal-to-noise properties of neural networks (Servan-Schreiber et al., 1990; Shine et al., 2021). Consequently, age-related decreases in dopaminergic neuromodulation, modelled as reduced neural gain, have been proposed to impoverish the distinctiveness of neural representations and networks (Li et al., 2001). Aligning with these computational accounts, we here demonstrate reduced D1DR availability to be associated with less specific functional coupling between the caudate and associative cortex across the adult lifespan. While caudate D1DR availability predicted differentiation of caudate-cortical connectivity during rest, a similar relationship was observed for prefrontal D1DR availability during working memory. The prefrontal D1DR-connectivity relationship during working memory persisted even after controlling for differences in caudate D1DR availability. This underscores the specific role of prefrontal dopamine in top-down regulation of cortico-striatal function during goal-oriented behaviour (Ott & Nieder, 2019), aligning with animal work indicating critical involvement of prefrontal D1DRs in working memory processes (Sawaguchi & Goldman-Rakic, 1991). Moreover, our work aligns with models postulating activation of D1DRs to promote the stability of networks patterns of activity (Durstewitz et al., 2000). Interestingly, previous work did not find striatal DAT availability to be associated with dedifferentiation of caudate-cortical connectivity in a smaller sample of older adults (Rieckmann et al., 2018), potentially suggesting a specific involvement of D1DRs. However, multi-tracer studies assessing pre- and post-synaptic components of the dopaminergic system within the same individuals are needed to evaluate this proposal.

Mediation analyses further indicated D1DR availability to account for age-related dedifferentiation of caudate-cortical coupling during rest and working memory, although we note that these results should be interpreted with caution due to the cross-sectional nature of the current data. While the present study primarily investigated dopaminergic regulation within specific cortico-striatal circuits, recent evidence also links striatal D1DR availability to functional organization of brain large-scale networks (Pedersen et al., 2023), suggesting potential brain-wide influences of striatal dopaminergic signalling beyond local circuits (McCutcheon et al., 2021). Moreover, we recently discovered that D1DR co-expression across the cortex follows a unimodal-transmodal hierarchy, exhibiting strong spatial correspondence to the principal gradient of functional connectivity (Pedersen et al., 2024). Taken together, our findings, along with these recent studies, suggest a tight coupling between D1DRs and functional connectivity at regional, network, and organizational scales.

Examining behavioural consequences of individual differences in caudate-cortical connectivity, we observed dedifferentiation of caudate-cortical coupling to predict poorer short and long-term memory performance in midlife and older age. This finding adds to evidence indicating decreasing functional and structural integrity of striato-cortical networks to be a key determinant of age-related deficits in flexible, higher-order cognition (e.g., Bonifazi et al., 2018; Fjell et al., 2016; Huang et al., 2017; Klostermann et al., 2012; Podell et al., 2012; Webb et al., 2020). Resting-state connectivity predicted performance on a battery of working and episodic memory tasks performed outside of the scanner, whereas connectivity during the working memory task displayed a similar relationship with in-scanner memory performance during the most challenging 3-back condition. The connectivity-behaviour associations observed across working and episodic memory tasks here likely reflect shared demands of these tests on executive control processes reliant on striatal and fronto-parietal circuits (Assem et al., 2020; Cools, 2019; Cools & D’Esposito, 2011; Zanto & Gazzaley, 2013). In contrast to the pattern of stronger connectivity-memory associations with older age observed in the current study, previous studies focusing on task activation have largely reported age-invariant associations between measures of neural (de)differentiation and cognition (reviewed in Koen & Rugg, 2019). It is possible that the lack of age moderation detected in previous studies may reflect modest sample sizes and low power for detection of such effects, or differences between the types of measures of neural specificity examined (i.e., regional activity vs. inter-regional functional connectivity).

While the current results highlight decreased dopaminergic integrity as one factor contributing to loss of specificity of neural function in older age, we acknowledge that other factors likely also play an important role. Animal (Leventhal et al., 2003) and human work (Chamberlain et al., 2021; Lalwani et al., 2019) implicate age-related decreases in inhibitory GABAergic neurotransmission in reduced distinctiveness of neural processing in aging, including decreased segregation of large-scale functional networks (Cassady et al., 2019). Similarly, declining integrity of white matter tracts has been shown to partly account for age-related changes in brain functional network organization (Pedersen et al., 2021). Fronto-striatal white matter connections are sensitive to age-related degradation (Webb et al., 2020), potentially contributing to the pattern of functional differences observed here. Moreover, while age-related decreases in the specificity of brain function have been reported across various metrics (e.g., univariate and multivariate patterns of regional activity, inter-regional functional connectivity), it remains to be elucidated whether shared or distinct mechanisms underpin age-related differences across these measures. Emerging evidence indicates correlations across representational and network levels of neural specificity (Cassady et al., 2020; Pauley et al., 2024), suggesting at least partly overlapping mechanisms, as predicted by neuromodulatory accounts (Li et al., 2001; Li & Sikström, 2002). Indeed, a recent study also demonstrated administration of the dopamine precursor L-DOPA to enhance neural representations underlying spatial navigation in young and older adults (Koch et al., 2022).

Finally, it is important to acknowledge that, in the present study, we examined caudate functional organization through the lens of caudate-cortical connectivity, utilizing a voxel-wise parcellation approach supplemented by ROI-based analyses. Although this approach has proven effective in characterizing striatal organization (e.g., Choi et al., 2012; Rieckmann et al., 2018), it necessitates imposing hard cut-offs between different subregions, assumes homogeneity within these subregions, and may overlook overlapping modes of functional organization (Haak et al., 2018; Marquand et al., 2017; O’Rawe & Leung, 2022). Future work could benefit from a gradient mapping approach to investigate overlapping modes of striatal organization, akin to our recent work on hippocampal organization (Nordin et al., 2024).

To summarize, we demonstrate advancing adult age to be associated with gradual dedifferentiation of caudate functional organization. Consistent with computational accounts of dopaminergic influences on age-related neural dedifferentiation, we offer novel empirical evidence identifying decreased D1DR availability in the striatum and the prefrontal cortex as predictors of less differentiated caudate-cortical functional connectivity across the adult lifespan. Less differentiated caudate-cortical connectivity was further associated with poorer memory performance in midlife and older age, underscoring the importance of functional integrity of cortico-striatal circuits for maintenance of memory abilities with advancing age.

