## Supplementary material for "Dedifferentiation of caudate functional organization is linked to reduced D1 dopamine receptor availability and poorer memory function in aging"

*Principal component analysis of memory performance*

*Table S1.* Loadings of individual episodic and working memory tasks on the first principal component that accounted for 54.96% variance in memory performance.

| **Task** | **Loading** |
|---|---|
| Word recall | .80 |
| Number-word recall | .64 |
| Object-location recall | .70 |
| Letter updating | .76 |
| Numerical 3-back | .77 |
| Spatial updating | .77 |

*Voxel-wise parcellations of resting-state caudate-cortical connectivity*

*Table S2.* Percentage of caudate voxels allocated to each cortical network in each age group, based on a 7-network parcellation (Yeo et al., 2011).

| **Network** | **Young** | **Middle-aged** | **Old** |
|---|---|---|---|
| Visual network | 2.71% | 1.70% | 2.91% |
| Somato-motor network | 0% | 0% | 1.30% |
| Dorsal attention network | 0% | 0% | 1.80% |
| Ventral attention network | 2.10% | 3.71% | 4.81% |
| Limbic network | 31.76% | 34.57% | 33.97% |
| Fronto-parietal network | 39.58% | 14.13% | 10.22% |
| Default-mode network | 23.85% | 45.89% | 44.99% |

*Controlling for educational level in the analysis of functional connectivity and memory relationships*

Including educational level as an additional covariate in the analyses of the relationship between functional connectivity and memory, we similarly observed connectivity differentiation during rest to predict out-of-scanner memory performance across the sample,

$\beta$ = .14, *SE* = .06, *t* = 2.40, *p* = .018, however, the interaction between age and connectivity became non-significant, $\beta$ = .08, *SE* = .05, *t* = 1.43, *p* = .154. Within the age groups, no significant association between connectivity differentiation and memory was still observed in the younger adults, $\beta$ = .04, *SE* = .13, *t* = 0.27, *p* = .790, whereas greater differentiation of caudate-cortical connectivity was positively associated with memory performance in the middle-aged, $\beta$ = .30, *SE* = .13, *t* = 2.39, *p* = .021, and older adults, $\beta$ = .34, *SE* = .14, *t* = 2.45, *p* = .018, even after controlling for education.

For the in-scanner n-back task, a significant 3-way interaction between age, memory load, and connectivity differentiation was similarly observed when including education as an additional covariate in the model, *F*(2, 341.86) = 3.53, *p* = .030. However, the within-group associations between connectivity differentiation and memory during the 3-back condition became non-significant in the middle-aged, $\beta$ = .21, *SE* = .13, *t* = 1.63, *p* = .110, and older adults, $\beta$ = .32, *SE* = .17, *t* = 1.84, *p* = .074, when additionally controlling for educational level.
